# A Novel Chloroplast Super-Complex Consisting of the ATP Synthase and Photosystem I Reaction Center

**DOI:** 10.1101/2019.12.25.888495

**Authors:** Satarupa Bhaduri, Sandeep K Singh, Whitaker Cohn, S. Saif Hasan, Julian P. Whitelegge, William A. Cramer

## Abstract

Several ‘super-complexes’ of individual hetero-oligomeric membrane protein complexes, whose function is to facilitate intra-membrane electron and proton transfer and harvesting of light energy, have been previously characterized in the mitochondrial cristae and chloroplast thylakoid membranes. The latter membrane is reported here to also be the location of an intra-membrane super-complex which is dominated by the ATP-synthase and photosystem I (PSI) reaction-center complexes, defined by mass spectrometry, clear-native PAGE and Western Blot analyses. This is the first documented presence of ATP synthase in a super-complex with the PSI reaction-center located in the non-appressed stromal domain of the thylakoid membrane.

## INTRODUCTION

Super-complexes of hetero-oligomeric electron transport membrane protein complexes, formed in the fluid mosaic membrane environment [1,2] have been described in mitochondria [3–6] where their formation has been shown to be facilitated by anionic lipid [3, 7–12]. The chloroplast thylakoid membrane contains four large hetero-oligomeric energy-transducing protein complexes (**Fig. 1**),for which molecular structure information has been obtained: (i) the photosystem II (PSII) reaction-center complex, responsible for the primary photochemistry associated with water splitting [13]; (ii) the cytochrome *b*_6_*f* complex [14] which provides the quinol acceptor, electron transfer connection, and trans-membrane proton translocation pathway [15, 16] between the PSII and photosystem I (PSI) reaction centers; (iii) the 561 kDa plant photosystem I (PSI) complex [17, 18] which is responsible for reduction of ferredoxin in the pathway of cyclic electron transport and reduction of NADP^+^ utilized for CO_2_ fixation; and (iv) the 560 kDa ATPase/synthase [19,20]. No super-complex has previously been documented which consists of complexes located in the non-appressed thylakoid region.

**Fig 1.**
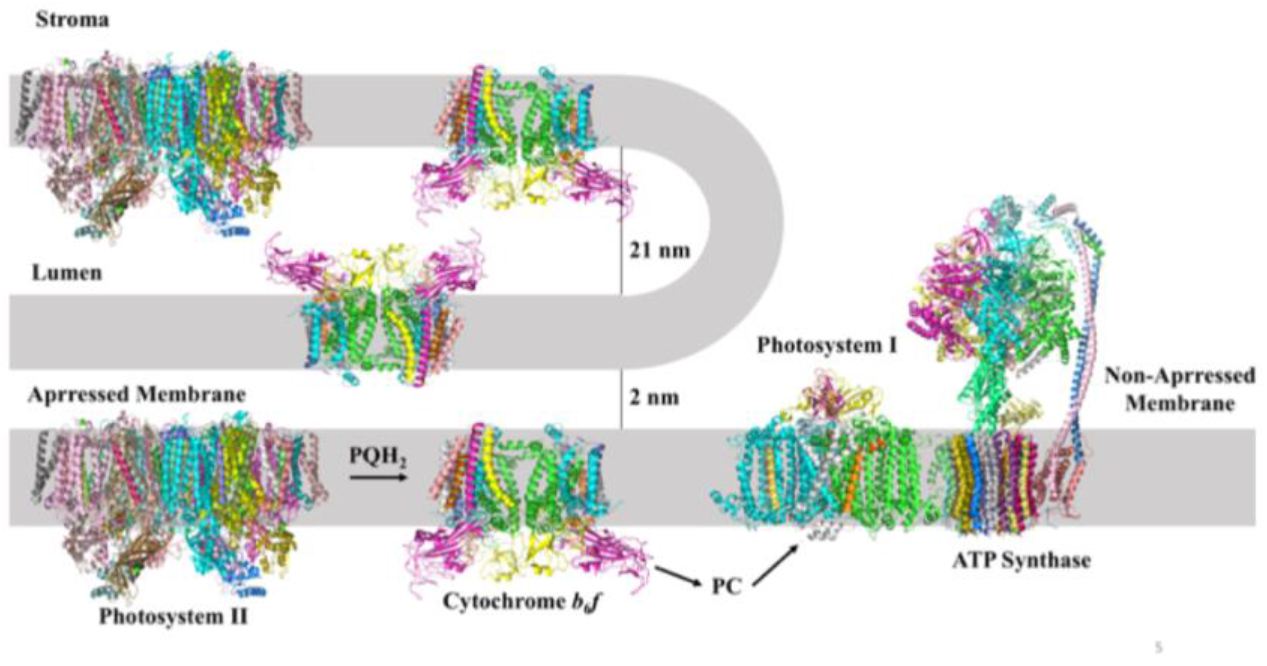
Distribution of the major intra-membrane energy transducing complexes associated with oxygenic photosynthesis in the appressed (grana) and non-appressed (stroma) thylakoid membranes. Photosystem II (PSII), photosystem I (PSI), cytochrome *b*_6_*f* complex, and ATP synthase, with molecular weights of approximately 350, 530, 270 (with FNR), and 560 kDa, respectively. A supercomplex of ATP synthase and photosystem I is implied in the figure by their proximity. Inter-membrane distances are obtained from the references [32, 41].

*Bipartite Thylakoid Membrane Structure; Residence of Energy-Transducing Complexes.* The chloroplast thylakoid membrane contains the energy transducing light-harvesting and electron-transport protein complexes arranged in two structurally distinct membrane domains, (i) the appressed grana membranes, and (ii) the non-appressed stroma or stroma-facing membrane domain (Fig. 1). The membrane locations of the PSI, PSII, *b*_6_*f*, and ATPase protein complexes have been discussed in the context of long-range redox communication [21], and the locations in grana and stromal compartments of the redox proteins that function in the ‘non-cyclic’ electron transport pathway from water to NADP^+^, and the PSI-linked redox proteins responsible for cyclic electron transport and linked phosphorylation, described [21]. The discovery of super-complexes in thylakoid membranes, which include complexes of the reaction centers with associated light harvesting chlorophyll proteins [22–30] and/or the *b*_6_*f* complex [31–33] and the electron transport/water-splitting structure [13] has led to an improved perspective on the organization of the membrane protein complexes which support the light harvesting and electron transport-proton translocation functions central to photosynthetic membrane energy transduction.

Super-complexes isolated from thylakoid membranes thus far are: (**i**) LHCII: PSII reaction center complexes [22–28]; (**ii**) nano-domains surrounding the PSII reaction center described by atomic force microscopy which are populated by the *b*_6_*f* complex [34] present in appressed and non-appressed membrane domains [32]; (**iii**) cytochrome *b*_6_*f* complex from *C. reinhardtii,* to which is bound light harvesting complexes LHCI and LHCII and ferredoxin-NADP^+^ reductase (FNR), which was obtained under conditions (‘state 1-state 2 transition’) favoring light energy transfer from the antenna pigment proteins to the PSI reaction center complex [32,33]; (**iv**) the *b*_6_*f* complex with PSI reaction center complex in *Arabidopsis* and *Chlamydomonas* [32,33]; (**v**) a super-complex of the PSI reaction center and the cytochrome *b*_6_*f* complex isolated from cold-adapted *C. reinhardtii* [35]; (**vi**) the structures described of the PSI reaction center complex and in association with LHCs from plant sources [29,30]; (**vii**) NDH-associated supercomplexes [32, 36]. The presence of the PSI reaction center and ATPase in the edge of the unstacked grana membranes has been documented [37–39].

The present study concerns a super-complex that combines the two hetero-oligomeric membrane protein complexes of the non-appressed thylakoid domain, ATP synthase and PSI reaction center, the first membrane protein super-complex in which ATP synthase has a major presence. This result has been documented by the use of high-resolution Orbitrap mass spectrometry to analyze the contribution to the super-complex of multiple major and minor protein components, and clear-native-PAGE, SDS-PAGE, and immuno-blot analysis to identify protein subunits of the component complexes.

## MATERIALS AND METHODS

### Purification of (i) Un-crosslinked Super-Complex (Preparations 1 and 2)

Spinach leaves macerated in darkness in HEPES-KOH pH 7.5, 150 mM NaCl, 5 mM EDTA at 4 °C and sedimented at 4000 x g, were re-suspended in HEPES (4-(2-hydroxyethyl)-1-piperazine-ethanesulfonic acid-KOH pH 7.5, 15 mM NaCl, 5 mM EDTA, centrifuged at 4,000 x g, and re-suspended in the same medium containing 5 mM NaCl. The chlorophyll concentration of thylakoid membranes was adjusted to 1 mg/ ml, extracted with 1 % DDM, centrifuged at 15,000 x g, the supernatant loaded on a 4-40% sucrose density gradient, and centrifuged at 200,000 x g.

### Purification of (ii) Crosslinked Super-Complex (Preparation 3)

Spinach leaves macerated in darkness in 50 mM Tris-HCl, pH 7.5, 0.1M NaCl, 0.3 M sucrose, and 1 mM EDTA, at 4 °C and sedimented at 6000 x g, were re-suspended in Tris-HCl, pH 7.5, 0.1 M NaCl, and 1 mM EDTA. The suspension was lysed for a second time using the French Press at a pressure of 8000 psi. The lysate was centrifuged at 10,000 x g, followed by ultracentrifugation at 160,000 x g for 30 min to separate stacked and unstacked grana thylakoids. The fractions were washed with 1 mM EDTA before the chlorophyll concentration was measured and adjusted to 2.5 mg/ml (chlorophyll a/b ratio = 5.4). Membrane proteins were extracted with 1 % β-DDM (β-dodecyl-maltoside), centrifuged at 17,000 x g, the supernatant loaded on a 4-40% sucrose density gradient, and centrifuged at 200,000 x g. The samples were crosslinked by 0.2% glutaraldehyde in the 40% sucrose solution used to make the 10-40% gradient.

### Electrophoretic Analysis

Protein bands from the sucrose gradient were fractionated, run on 4-12 % clear-native PAGE, and eight resolved complexes excised for mass spectrometric analysis. The largest band of approximately 1 MDa on the native gel (**Fig. 2B**) is inferred to contain the ATP synthase-PSI super-complex.

**Fig 2.**
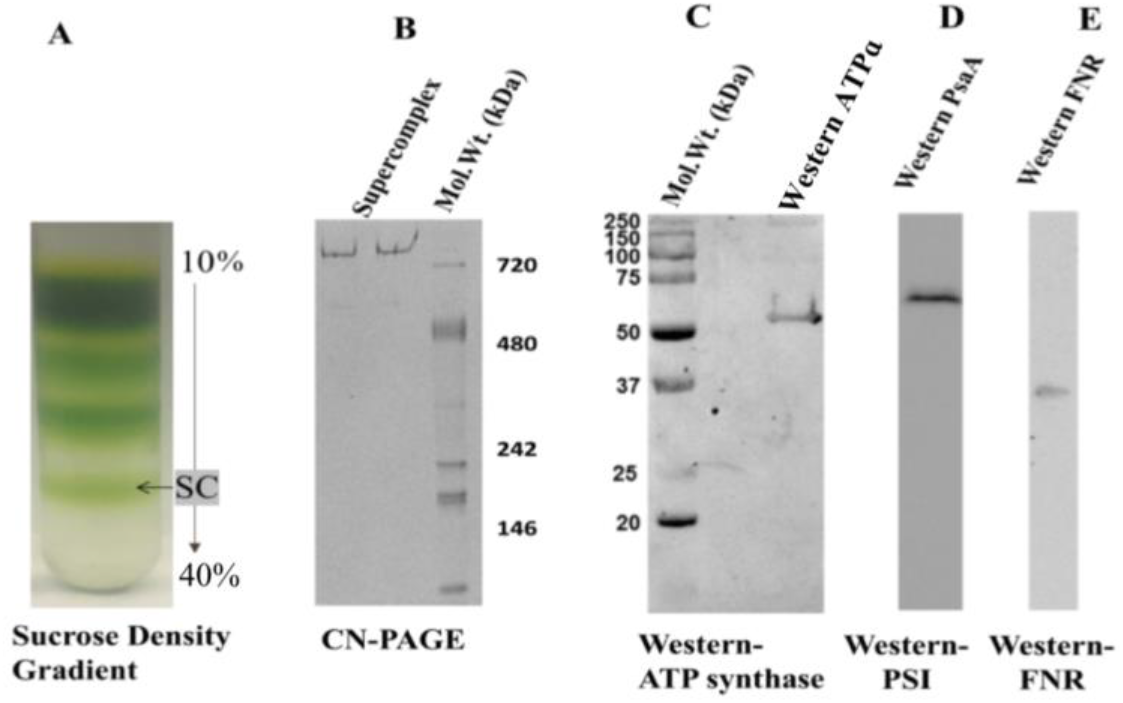
Purification and characterization of the PSI-ATP synthase super-complex. **(A)** 4-40 % sucrose density gradient of spinach thylakoid membrane extract. Super-complex (SC), marked by the arrow, is separated for further characterization. **(B)** 4-12 % Clear Native PAGE of the super-complex fraction isolated from the sucrose gradient and stained with Coomassie Brilliant Blue R-250. (**C, D, E**) Western blots of super-complex using 12% SDS-PAGE probed with antibodies raised against (C) α-subunit (55 kDa) of **Arabidopsi** ATP synthase, (D) PsaA subunit (60 kDa) of PSI, and (E) ferredoxin-NADPH oxidoreductase (FNR).

### Western Blotting

Isolated super-complex was assayed on SDS (data not shown) and clear-native PAGE, and transferred to a nitrocellulose membrane, the blot probed with anti-PsaA of PSI, anti-ATP synthase α-subunit, and anti-FNR antibodies obtained from PhytoAb, Inc. (**Fig. 2C,D and E**).

### Mass Spectroscopy

Mass spectrometry was performed as described previously using nano-liquid chromatography and a bench top Orbitrap mass spectrometer (QE-Plus, Thermo-Fisher Scientific) operated in positive ion mode (nLC-MSMS) [40]. Proteins in native gel-bands were alkylated, digested with trypsin and extracted for nLC-MSMS. The Mascot Search Algorithm (Matrix Science, London, UK) was utilized to identify the significant presence of peptides, using 10 ppm mass tolerance on high-resolution precursor MS1 scans, and 20 milli-mass unit tolerance on high-resolution product MS2 scans, obtaining an overall < 5% peptide false discovery rate based upon a reversed sequence decoy search [41]. A spinach database was downloaded from Uniprot (UP000054095). The top twenty Mascot protein scores for each sample were presented representing the most abundant proteins in the sample and minimizing low scoring proteins that are more likely to represent noise (Mascot ‘protein’ score is the summed score for unique peptides within the sample analysis). A ‘complex’ score for each of the four major thylakoid protein complexes was generated by adding together the protein scores for the top four highest scoring proteins (or fewer than four if there were fewer than four proteins from that complex in the twenty highest scoring proteins). The top twenty proteins were chosen as a cutoff because proteins below this threshold have protein scores around 10% or less of the highest scoring proteins and thus are more likely to be in the ‘noise’ fraction [32].

### Assay of ATPase Activity

The “ADP-Glo” kit (Promega, Wisconsin) was used to measure hydrolysis of ATP to ADP and inorganic phosphate by ATPase. ATP (0.4 mM) was added to the super-complex fraction in the presence of buffer (50 mM Tris-HCl, 5 mM Mg^2+^, and BSA) at room temperature, incubated for 15 min, followed by the addition of “ADP-Glo reagent” and incubation at room temperature (40 min) to consume unused ATP. The reaction mixture was then incubated with the ‘detection reagent’ to convert ADP to ATP, measured by the reaction of luciferase/luciferin, and the luminescence recorded in a spectral band centered at 560 nm, utilizing a Spectramax Luminometer (Molecular Devices, CA).

### Fluorescence Emission Spectroscopy

Fluorescence measurements were made with a FluoroMax 3 Fluorimeter (Horriba-Yvon Inc.), using a 3 x 3 mm quartz microcell with 0.3 ml volume. FAD containing the super-complex fraction in 10 mM HEPES buffer, pH 7.5, 0.5 mM EDTA, 2 mM DDM, 10 % sucrose. Fluorescence was excited at 460 nm, and emission spectra collected in a 500-580 nm spectral interval, with a stepping increment of 1 nm and a 0.1 sec integration time. Excitation spectra were measured from 420 – 500 nm using an emission band interval centered at 530 nm. Difference spectra were obtained by measuring the fluorescence in the presence and absence of sodium dithionite, the latter a reductant with a redox potential negative enough to fully reduce the flavin in FNR, thus effectively eliminating its absorbance and resulting fluorescence.

## RESULTS

### A unique super-complex from thylakoid membranes; electrophoretic and immuno-blot analysis

The super-complex was isolated from spinach thylakoid membranes by detergent extraction followed by sucrose density gradient centrifugation (**Fig. 2A**, marked as “SC”), and the approximate size was determined on a 4–12 % clear-native gel (**Fig. 2B**).

The presence of ATP synthase, PS I and FNR was confirmed by Western blot analyses using antibodies raised against the *Arabidopsis* ATP synthase α-subunit (55 kDa) (**Fig. 2C**), the 60 kDa PsaA subunit of PSI (**Fig. 2D**), and FNR (36 kDa) (**Fig. 2E**).

### Mass Spectrometry

High-resolution mass spectrometry of the isolated 1 MDa super-complex from spinach chloroplast thylakoid membranes (**Table 1**) revealed the presence of ATP synthase (0.56 MDa), Photosystem I (0.53 MDa). Polypeptide components derived from the 1MDa band in **Fig. 2B**, were identified after in-gel digestion of the native gel-band and tandem mass spectrometry with database searching constrained by a specified high statistical confidence threshold, as described in *Methods*, and summarized in **Table 1A, B**. Presence of ATP synthase and Photosystem I were identified with high statistical confidence by multiple polypeptides. Notably absent were Photosystem II (trace amounts of PsbB were detected in 1 preparation, assumed to represent contamination), and cytochrome *b*_*6*_*f* complex polypeptides. Other components that were detected consistently were the polypeptides FtsH and FtsH2, Lhca3.1 (a known PSI chlorophyll a/b binding protein) and ferredoxin-NADPH oxidoreductase (FNR) that was confirmed by Western blot (**Fig. 2E**). A calcium-sensing receptor was found in 2/3 preparations and rubisco activase was found with a high score after crosslinking. Because of differential recovery of larger versus smaller subunits and peptides derived from them it is not possible to infer polypeptide stoichiometry from these measurements.

**Table 1A.**
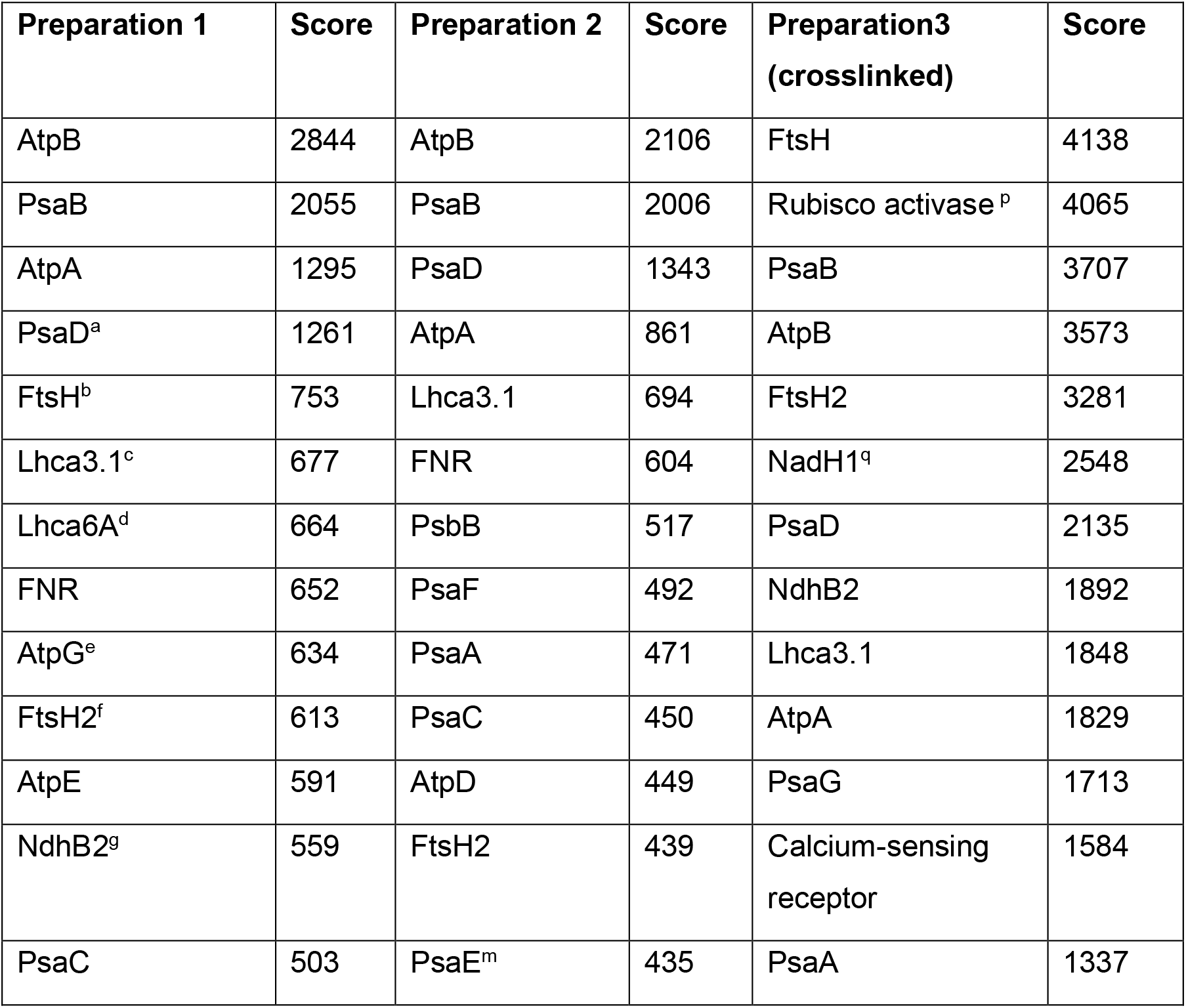
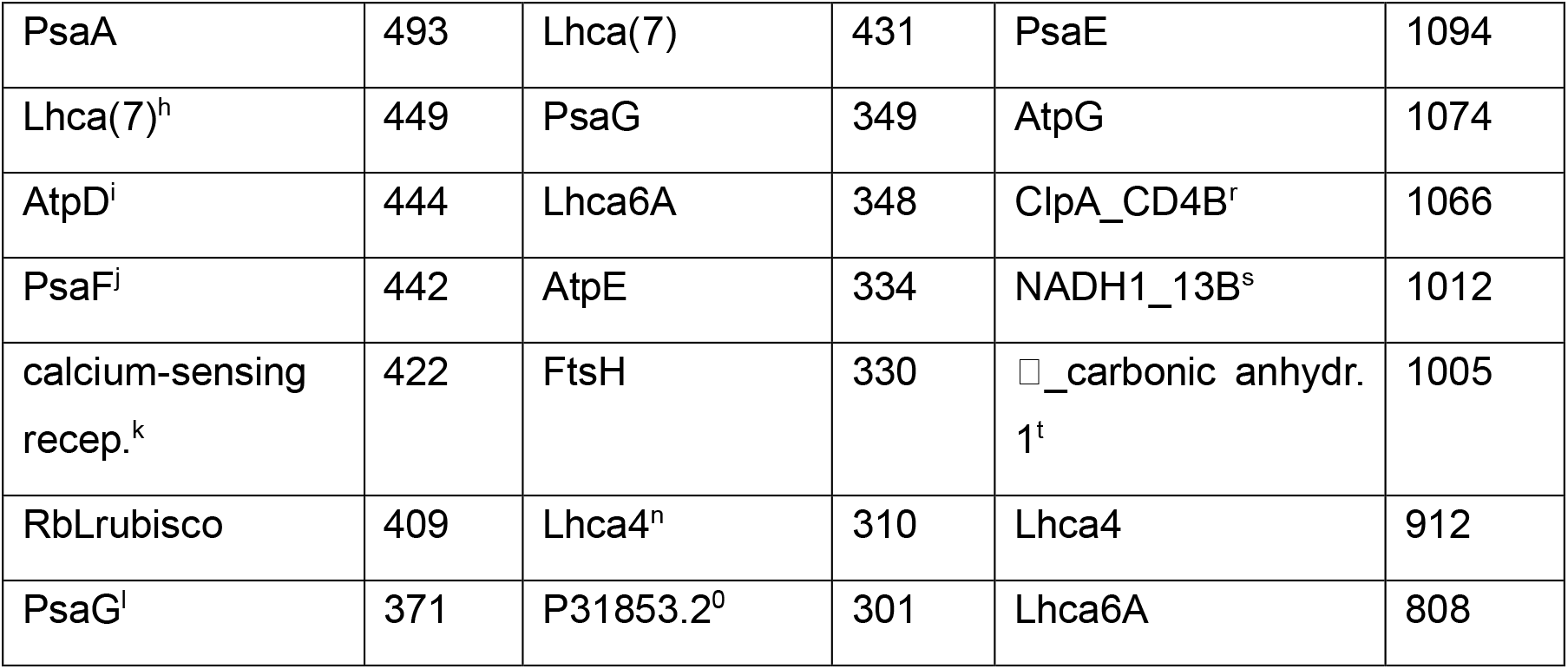
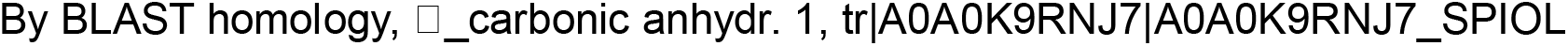
Identification of Twenty Highest Scoring Proteins in Three Super-Complex Preparations.

**Table 1B.** Major Thylakoid Complexes in Supercomplex.

| Complex | Prep. 1 | Prep.2 | Prep.3<br>(crosslinked) |
| --- | --- | --- | --- |
| ATPase | 5364 | 3750 | 6476 |
| PSI | 4312 | 4312 | 8892 |
| $b_6f$ | 0 | 0 | 0 |
| PSII* | 0 | 517<sup>c</sup> | 0 |

\*Minor contamination in by PSII.

4 highest scores for each complex component added together.

The presence of FNR was confirmed through Western blot analysis using an anti-FNR antibody probe (**Fig. 2E**), and determination of fluorescence excitation and emission spectra that are characteristic of FAD [42] (**Fig. 3**). No polypeptides from the cytochrome *b
_6_f* complex were detected at the stringency used.

**Figure 3.**
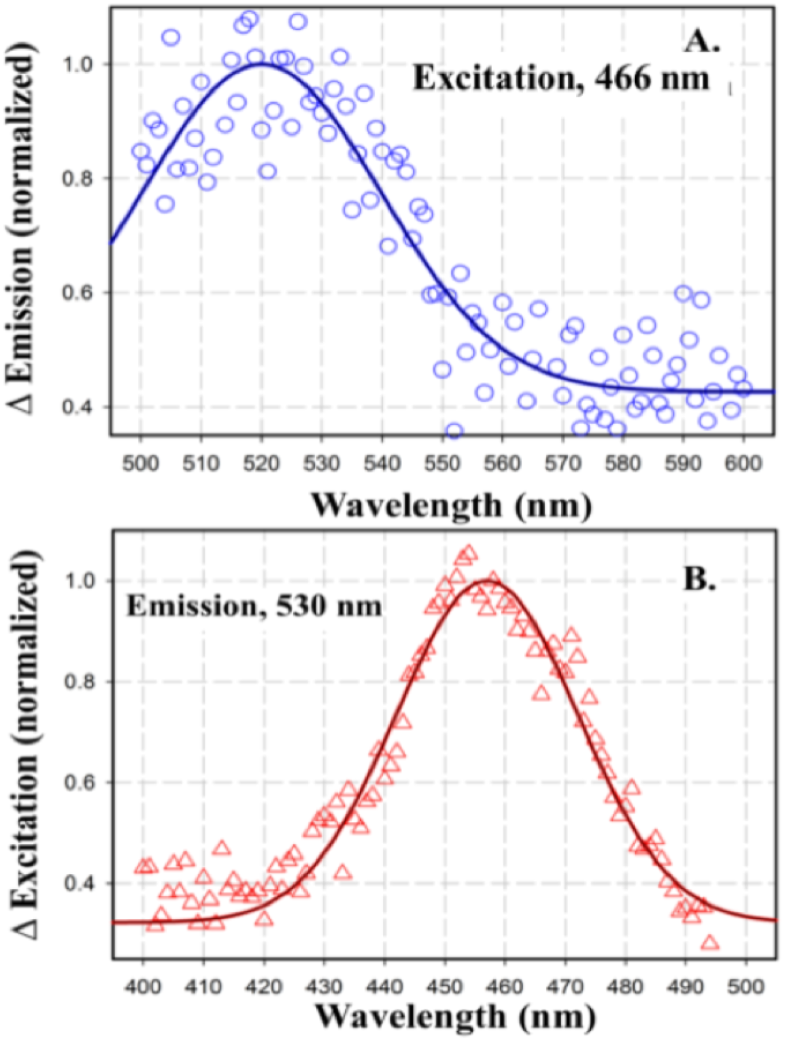
Difference fluorescence emission (top) and excitation (bottom) spectra of the ‘super-complex;’ measured with a FluoroMax 3 Fluorimeter (Horiba-Yvon Inc.). **(A)** Emission spectra measured between 500 – 600 nm. Stepping increment, 1 nm; Integration time, 0.1 sec; Excitation, 466 nm; bandwidth 5 nm for excitation and emission. **(B)** Excitation spectra measured in 400–500 nm range, measuring the emission at its 530 nm peak. Spectra of undiluted sample were measured in a quartz microcell, 3 x 3 mm, volume, 0.3 ml (Starna Cells) before and after addition of dithionite grains. Difference spectra were obtained by subtraction of spectra, i. e., ‘oxidized–reduced,’ measured before and after addition of dithionite.

### Presence of FNR in the Super-Complex

The presence of FNR in the spinach super complex was confirmed from fluorescence emission and excitation spectra produced by the isoalloxazine moiety of FAD (**Figs. 3A, B**). The super-complex, suspended in 10 mM HEPES buffer, pH 7.5, and 0.5 mM EDTA, 2 mM DDM, 10 % sucrose, was excited at 460 nm and emission spectra were acquired over the wavelength range 500-600 nm. The difference emission spectrum, calculated with and without the strong reductant, sodium dithionite, showed an emission maximum in the oxidized state at 530 nm, which is characteristic of the presence of flavin [42]. A similar conclusion was reached from the difference excitation spectra measured with and without sodium dithionite over the 400-500 nm range using emission at 530 nm. The excitation spectrum showed maximum at 460 nm, characteristic of oxidized flavin [42]. Spectra are normalized to unit emission intensity.

### ATPase Activity

The F1-ATPase component in the super-complex demonstrated *in vitro* ATP hydrolysis, as described in *Methods*. The “ADP-Glo kit” (Promega) was used to provide a qualitative measure of ATP hydrolysis, which was dependent on the presence of the PSI-ATPase-FNR super-complex (**Table 2**). ATP hydrolysis was measured by addition of 0.4 mM ATP to the super-complex fraction in the buffer containing 50 mM Tris-HCl, 5 mM Mg^2+^, and 0.1 mg/ml BSA).

**Table 2.** ATPase Activity of the Super-Complex. As described in *Methods*, kinase activity was measured qualitatively through luciferase emission in the positive control (buffer and 0.4 mM ADP); negative control (buffer and ATP (0.4 mM) and the purified super-complex in buffer and 0.4 mM ATP.

| Sample | Chemiluminescent Signal (cpm) |
| --- | --- |
| +ADP (positive control) | $(3.1 \pm 2) \times 10^6$ |
| +ATP (negative control) | $(5.5 \pm 1.2) \times 10^2$ |
| + Supercomplex | $(4.4 \pm 0.6) \times 10^5$ |

## DISCUSSION

Several super-complexes of PSI with thylakoid complexes have previously been described [17,18, 29, 30], but a complex with ATP synthase has thus far been absent. These authors defined the super-complex based upon the ability to visualize the complex using cryo-electron microscopy. In the present case it has not been possible to preserve the PSI-ATP synthase complex during the freezing process for cryo-EM. Consequently, we do not include an EM image of this super-complex in this study. Alternatively, classical biochemical techniques show this super-complex running as a discrete band in sucrose-gradient centrifugation and as a ~1 MDa band on a native gel combined with a modern description of the major components of the super-complex by high-resolution mass spectrometry.

The identity of the PSI-ATP synthase super-complex is supported by the facts that (i) the super-complex band is the densest, implying a higher protein/lipid content than other thylakoid super-complexes consistent with the presence of the ATP synthase which contains no pigments; (ii) the molecular weight observed by native gel electrophoresis of ~1 MDa is consistent with an equimolar mixture of PSI [17,18] (MW= 561 kDa) and ATP synthase (MW = 560 kDa) [19,20], and no other complex.

Modern mass spectrometry using high-resolution precursor-and production scans in tandem MS protein identification underscores current proteomics [44]. In-gel analysis of bands on native gels is relatively uncommon compared to bands on SDS gels. In fact, the number of tryptic peptides derived from the subunits of a super-complex, far exceeds that expected from trypsin treatment of a single protein. The modern mass spectrometer combined with reverse-phase chromatography provides a powerful tool to analyze a heterogenous mixture of peptides to determine the super-complex. Integral membrane proteins with large extrinsic domains yield abundant peptides that are easily recoverable in the in-gel procedure yielding very high relative abundance ‘scores’ where these subunits are present. Thus, ATP synthase and PSI with very high scores are identified, based on the four highest scoring polypeptides underlying our conclusion that these two complexes are the only complexes within this super-complex (**Table 1B**). PSII and cytochrome *b_6_f* components are easily identified in other bands from the sucrose gradient (not shown) but not in the densest band representing the super-complex as described. Small subunits with a large proportion of transmembrane sequence are poorly represented in this type of analysis [45] and were not seen in this analysis. Intact protein analysis is better suited for full subunit coverage, as reported in 2002 [46] and would be well suited to detect all the subunits in the supercomplex. Native intact supercomplex mass spectrometry would be most desirable as it addresses stoichiometry [47], but it as yet unavailable for this preparation.

Mass spectrometry also showed the presence of other proteins that might be harder to define by cryo-EM, reliably identifying FtsH and FtsH2, Lhca3A and FNR. Initial studies on these components have documented the presence of FNR.

This is the first instance of detection of a photosynthetic super-complex which contains the ~560 kDa ATP synthase protein complex. Localized coupling of the electron transport system and ATP synthesis by super-complex formation in mitochondria has been proposed [48]. It has been suggested that the oxidative phosphorylation system in respiratory systems can operate in two states: via utilization of (i) the bulk phase trans-membrane electrochemical potential gradient [48], or (ii) protons localized in the membrane that are accessible to the active site chemistry [49]. In the present study, the occurrence of ATP synthase in association with the PSI reaction-center complex, suggests the possibility of a relatively localized coupling between (i) ATP synthesis and the transmembrane electrical potential generated in the PSI reaction center, and/or (ii) a localized domain of carbon dioxide fixation.

The isolated complex dominated by the presence of PSI and ATP synthase is also associated with a number of proteins like the ATP dependent, membrane anchored zinc metalloprotease FtsH, FNR and Rubisco Activase. FtsH has been implicated in chloroplast biogenesis and repair of photosystem II under high light conditions [50]. It has been demonstrated that FtsH, localized in the stroma lamellae, is critical for the biogenesis of PSI complex under normal light conditions [51]. In bacteria FtsH is responsible for regulating ATP synthase biogenesis [52]. However, such a function has not yet been documented in the plant system.

Previous studies have shown that a Chl a/b ratio of 4.6 in the stroma lamellae isolated from spinach leaves, had a PSI/PSII ratio of ~3 [53]. In the current preparation, a chlorophyll a/b ratio of 5.4 was measured in the membrane fraction, which indicates that the super-complex PSI-ATPase containing has been derived from a PSI enriched stroma lamellae membrane. A high relative abundance of Lhca 3.1, a light harvesting protein associated with PSI [54], as determined by mass spectrometry, also confirms a major presence of photosystem I in the super-complex.

The flavoenzyme, ferredoxin-NADP^+^ reductase (FNR), found in both soluble and thylakoid membrane associated forms, has been proposed to define the photosynthetic electron transport pathway [55]. PSI bound FNR or soluble stromal FNR facilitates linear electron flow, while under reduced conditions, FNR associates with the cytochrome *b*_6_*f* complex and promotes cyclic electron flow [56]. The presence of FNR in the super-complex has been confirmed by mass spectrometry (**Table 1A**), Western Blot (**Fig. 2E**) and difference fluorescence spectroscopy (**Fig. 3**). A third protein, rubisco activase, found associated with the high molecular weight crosslinked band containing PSI and ATP synthase (Preparation 3), interacts with Rubisco to release the bound inhibitors in an ATP dependent fashion and hence, initiates the CO2 fixation cycle [57].

## AUTHOR INFORMATION

### Author Contributions

Project proposed by SKS and WAC. Super-complex isolation and biochemical characterization done by SKS and SB. Mass spectrometry analyses done by JPW. SSH contributed to the experimental design. Manuscript written by SB, JPW, and WAC.

## ABBREVIATIONS

CN: clear native
SDS-PAGE: sodium dodecyl sulfate-polyacrylamide gel electrophoresis
cpm: counts/min
cyt: cytochrome
DDM: n-dodecyl-β-D-malto-pyranoside
FNR: ferredoxin-NADP^+^ reductase
EM: electron microscopy
FtsH: filamentous, temperature-sensitive ATP-dependent protease
HRP: horse radish peroxidase
LHC: light harvesting complex
Mr: relative molecular weight
PAGE: polyacrylamide gel electrophoresis
pdb: protein data bank
PSI: PSII, photosystems I, II

## ACKNOWLEDGMENTS

We thank S. D. Zakharov, S. Puthiyaveetil, R. Morgan-Kiss, and I. Kalra for helpful discussions, and S. Naurin for technical support. Studies were supported by Dept. of Energy/Basic Energy Sciences/Photosynthetic Systems grant DE-SC00118238, the Henry Koffler Distinguished Professorship (WAC), and NIHDK-063491 (JPW).

